# Establishment of a closed artificial ecosystem to ensure human long-term survival on the moon

**DOI:** 10.1101/2021.01.12.426282

**Authors:** Yuming Fu, Zhihao Yi, Yao Du, Hui Liu, Beizhen Xie, Hong Liu

**Affiliations:** Beijing Advanced Innovation Centre for Biomedical Engineering, Beihang University, Beijing, 100083, China; School of Biological Science and Medical Engineering, Beihang University, Beijing, 100083, China; International Joint Research Center of Aerospace Biotechnology & Medical Engineering, Beihang University, Beijing 100083, China

**Author notes:** **Corresponding author:** Hong Liu, Ph.D, Professor, Beihang University.

**Keywords:** Ecosystem ecology, bioregenerative life support system (BLSS), lunar base, atmospheric stability, material circulation

## Abstract

Bioregenerative life support system (BLSS) is a critical technology maintaining long-term human survival on the Moon or other extraterrestrial bodies. In the current study, we carried out a 370-day integrated high-closure experiment (“Lunar Palace 365” experiment) on the Earth in an upgraded ground-based BLSS experimental facility called “Lunar Palace 1”. This experiment was designed to develop techniques to run and adjust system stability under long-term operation and crew shift change conditions. Eight volunteering crew members were divided into two groups with three time phases: Group I stayed in the cabin for the initial 60-day phase; Group II inhabited the cabin instead for a record-breaking duration of 200 days as the second phase; Group I re-entered the cabin, replaced Group II and stayed for the last 105 days. Our results demonstrated the BLSS had excellent stability with a material closure degree of 98.2%. Levels of O_2_, CO_2_ and trace harmful gases were well controlled within ranges optimal for crew health and plant production. The system exhibited a strong robustness and could quickly minimize effects of disturbances through self-feedback adjustments. The efficiency of plant production completely met the crew’s need of plant-based food. The purification efficiency of domestic and sanitary wastewater was up to irrigation standards, and the recovery rate of urine and solid waste achieved 99.7% and 67%, respectively. These results are valuable for further optimization of the BLSS in a lunar base and computer simulations of similar systems.

## 1. Introduction

Space science is developing vigorously, especially regarding manned landing on the Moon. Long-term lunar bases will be established in the near future to better explore the Moon. This requires for more verifications and studies, which also has significance of reference for further Martian explorations. It is a huge challenge to live outside the Earth and thus important to build Bioregenerative Life Support System (BLSS), closed artificial ecosystem based on ecological principles, in which oxygen, water, and major food sources are recycled using biotechnology and engineering control technology, creating an Earth-like microenvironment^1^. BLSS is the most advanced life support technology, but also the most complex and highest-priority technology in long-term manned deep space exploration activities such as future, long-term lunar bases^2^. Before advancing into practical space exploration, techniques running and adjusting a steady BLSS must be first developed through simulation experiments on the Earth. This will provide an experimental basis and a data reference for BLSS engineering applicable to lunar bases. To meet this, from the 1970s and ‘80s to the beginning of 21st century, several countries, including Russia (BIOS-3 system)^3^, the United States (BIO-Plex system)^4^, Japan (CEEF system)^5^ and the European Union (MELLISA system)^6^ have performed crewed simulation experiments of BLSS on the Earth.

In recent years, with the rapid development of manned spaceflight industry, breakthroughs in BLSS unit technologies, particularly the development of highly efficient plant cultivation^7–9^, animal protein production^10,11^, nitrogen recovery from urine^12^, and bioconversion of solid wastes into soil-like substrates^13–15^, have been achieved. Based on the development of BLSS unit technologies, teams from China have carried out research to integrate these technologies into BLSS. In previous studies, much attention was paid to the regeneration of O_2_ and mass flow in a BLSS. For instance, microalgae and lettuce had been shown as meeting the O_2_ demand of 1/2 of people in 297 days of closed experiments^16^; *Azolla* photosynthetic O_2_ release was able to achieve O_2_-CO_2_ homeostasis in the “*Azolla*-fish-men” closed system^8^, while a 13.5 m^2^ planting area met the O_2_ demand of one person in a two-person, 30-day “lettuce-human” integrated controlled ecological life support system (CELSS) test^17^. More recent studies investigated the regeneration rate of other BLSS components of longer durations and more diverse crew conditions. For example, in 2014, our group carried out the first 105-day integrated experiment in a ground-based BLSS facility, Lunar Palace 1^2,18^. In that experiment, we used straw to feed yellow mealworms to provide animal protein for volunteering crew members, achieving 100% recycling of O_2_ and water and regenerating most foods, with an overall closure degree of 97%. To explore the establishment of CELSS for future long-term extraterrestrial stays, a four-person and 180-day CELSS integrated experiment was performed by the Astronaut Research and Training Center of China in 2016 ^19^. That study systematically converted the material regeneration model from physical-chemical to biological regeneration, and confirmed the applicability and reliability of each unit’s technology. Despite the significant progress in BLSS integration research, there is still a need to establish a BLSS which will function well over long-term continuous operation in an extraterrestrial environment. In addition, when performing long-term space missions, such as exploration of the Moon, shift changes of crew members are inevitable. It is therefore important to understand how a BLSS can run steadily during shift changes which could largely affect the metabolic rate of the system. Impacts of power failures and other equipment malfunction during the operation should also be considered to develop a viable BLSS.

In the current study, we carried out a ground-based 370-day, integrated high-closure experiment (“Lunar Palace 365” experiment) to explore long-term life support needs of a lunar-base-like habitat with multiple crew member shift changes. Such an experiment will improve not only the system closure degree and stability in long-term operations, but also our knowledge of human life support regarding long-term Moon habitation.

## 2. Materials and methods

### 2.1 Lunar Palace 1, a ground-based artificial, closed ecological facility

The “Lunar Palace 365” program to test a life support system for 365 days was carried out in “Lunar Palace 1”, a ground-based comprehensive, experimental BLSS facility located in Beihang University (BUAA). Lunar Palace 1 was designed to contain two plant cabins and one comprehensive cabin (Fig. 1), with a total area of 160 m^2^ and volume of 500 m^3^. Each plant cabin was divided into two parts (plant cabin I and II) with independent environmental condition control. The plant cabin was critical to the regeneration of O_2_ and water as well as food production. The comprehensive cabin, the main living space for the volunteer crew members, hosted four one-bed cubicles, a bathroom, an animal-raising room, a living room, a storage room and a waste-treatment room. The whole system was closed and completely air-tight without any material interaction with the outside. The actual gas leakage rate of the cabin at normal pressure was only 0.043%/d^2,20^, except for three time points wherein intentionally simulated accidents were carried out. The temperature and humidity in plant cabin I during the whole experiment were 22.25±1°C and 62.09 ±5.28%, and 22.72±0.89°C and 61.52%±5.11% for plant cabin II. Temperature and humidity control of the plant cabin was stable, and conformed to typical planting conditions^2,3,21^. Temperature and humidity fluctuations of the comprehensive cabin were 24.11±1.36°C and 49.62 ±4.96%, respectively, which met the requirements of the advanced life support system habitation cabin (temperature: 18.5-26.5°C; relative humidity: 25%-70%) specified by NASA standards^21^.

**Fig.1.**
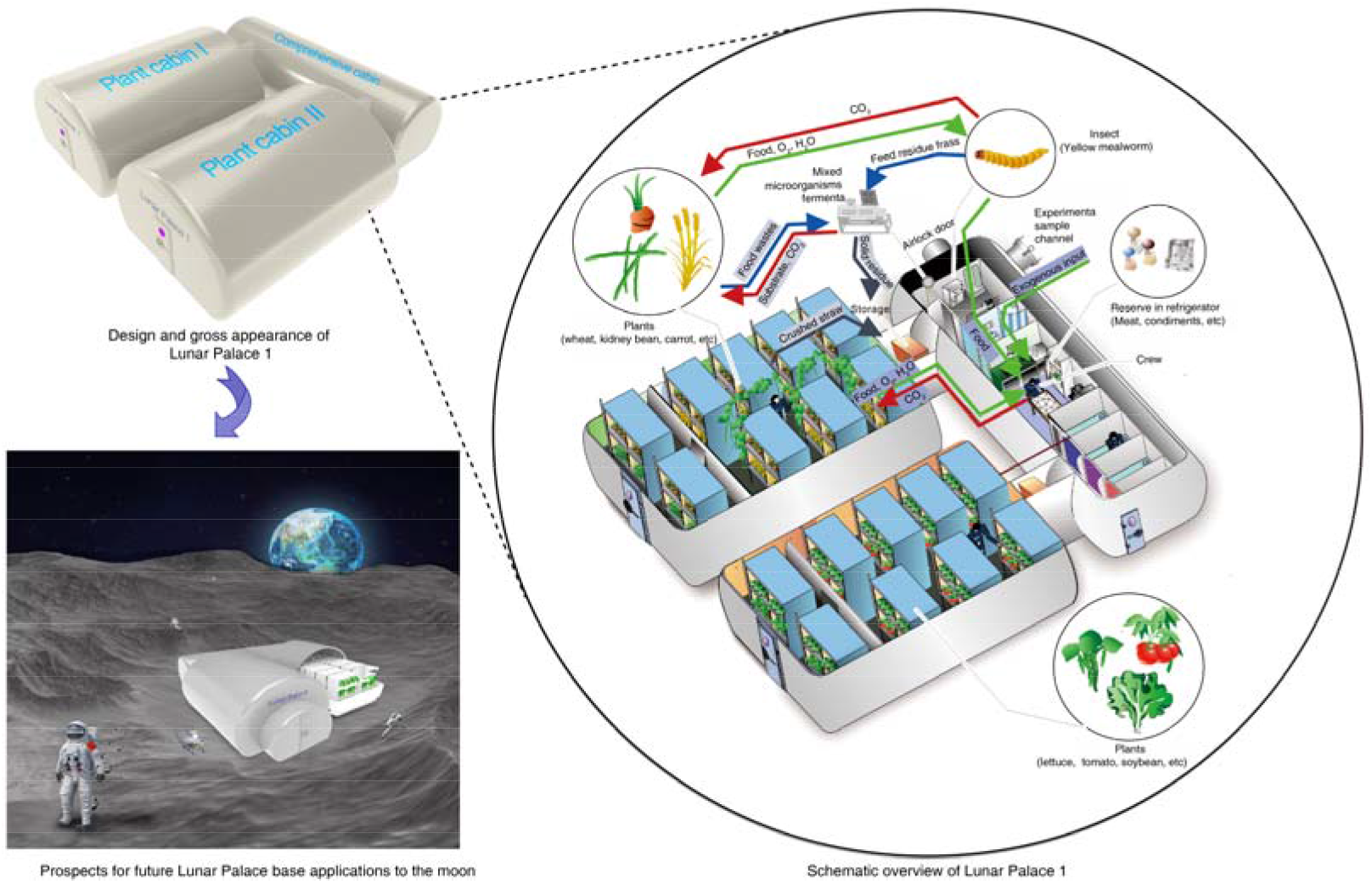
The layout, schematic and future application of Lunar Palace 1.

### 2.2 The biological regeneration cycle of Lunar Palace 1

Lunar Palace 1 was a BLSS composed of a “human-animal-plant-microorganism” ecosystem, wherein the crew members cultivated food plants, vegetables, and fruits as the crew’s food (Fig. 1). The plants absorbed CO_2_ from the air and generated O_2_ via photosynthesis, with the O_2_ used in turn for respiration by the crew, animals, and microorganisms. Thus, regeneration of O_2_ was realized in the BLSS system, maintaining the gas balance of the system.

Water output of the crew included domestic wastewater and urine (Fig. 2A). Domestic wastewater was transferred to a membrane biological activated carbon bioreactor (MBACR) for purification; the urine of the crew went into rotary evaporation to retrieve elemental nitrogen and water, which was subsequently transferred to the MBACR for purification too. The reclaimed water from the MBACR was transferred into plant nutrient solution preparation tank and then disinfected by UV for plant growth. The irrigation water was purified by evaporation and plant transpiration, and was transited to liquid water in plant cabin condensate. After MBACR purification and UV disinfection, the water was then ready for direct use of the crew, including drinking water, sanitary water and domestic water.

**Fig.2.**
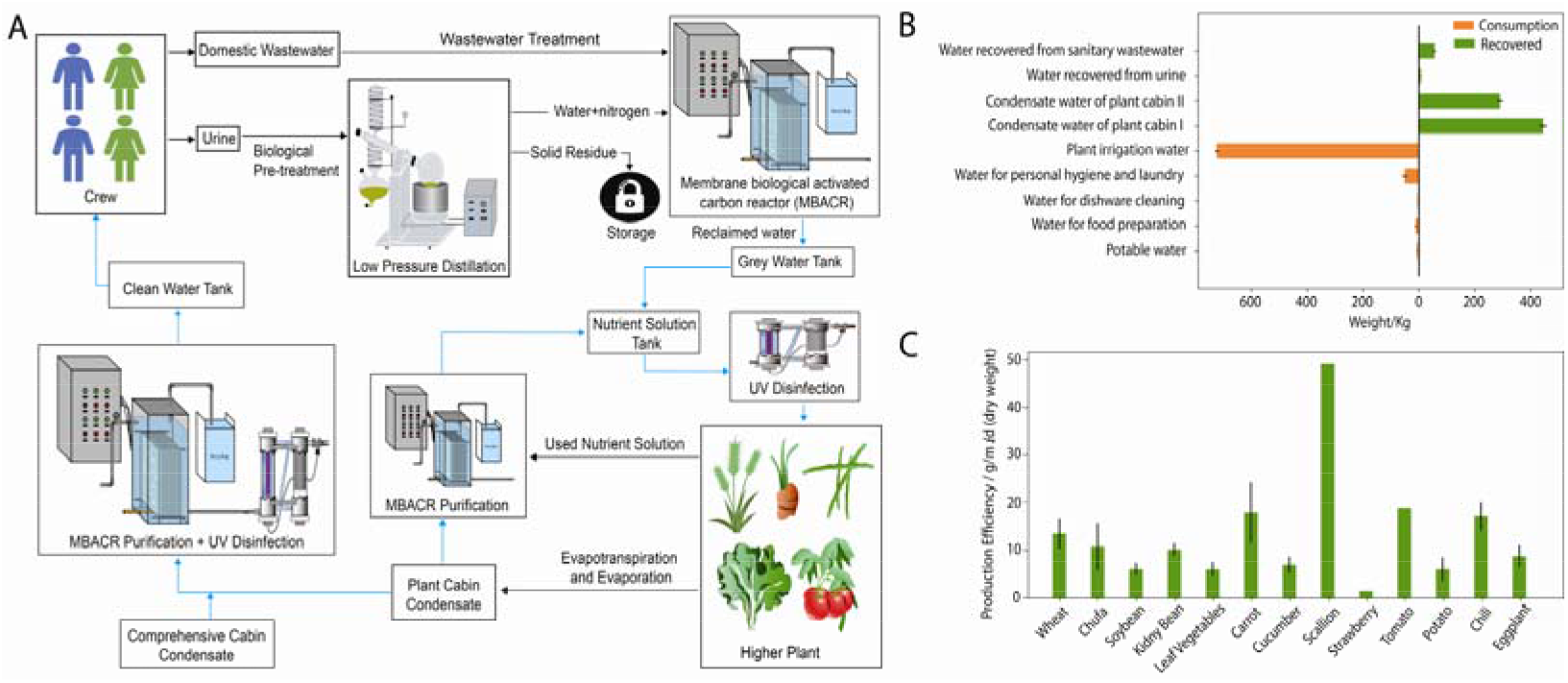
The water cycle and food Production. (A) BLSS water circulation and treatment processes in the “Lunar Palace 365” experiment. (B) Water consumption and recovery (kg/d) during the “Lunar Palace 365” experiment. (C) Production efficiency of 13 selected crops (g m^−2^ day^−1^, dry weight) in the “Lunar Palace 365” experiment.

The grain straw was dried and crushed after harvest, the better part of which were mixed with feces of the crew and placed into solid fermentation unit inoculated with a straw decomposition bacterial agent. A small amount of straw was fermented, and another small amount of straw was used to cultivate mushroom. The fermented straw and mushroom substrate were then mixed with the leaves of harvested vegetable plants as feed for insects (yellow mealworms). The feed residue (including insect dung) generated by the insects were also placed into the fermenter. Thus, solid wastes, including crew feces, insect feed residue and straw, were mixed and fermented in the solid fermentation unit. The CO_2_-rich gas generated by fermentation was purified by air purifiers and then introduced into the plant cabin for photosynthesis. Some of the solid products after fermentation were used for plant cultivation, while others were collected, compressed and stored. Yellow mealworms provided some animal protein for the crew. The process of solid waste recycling is shown in Fig. 3A.

**Fig.3.**
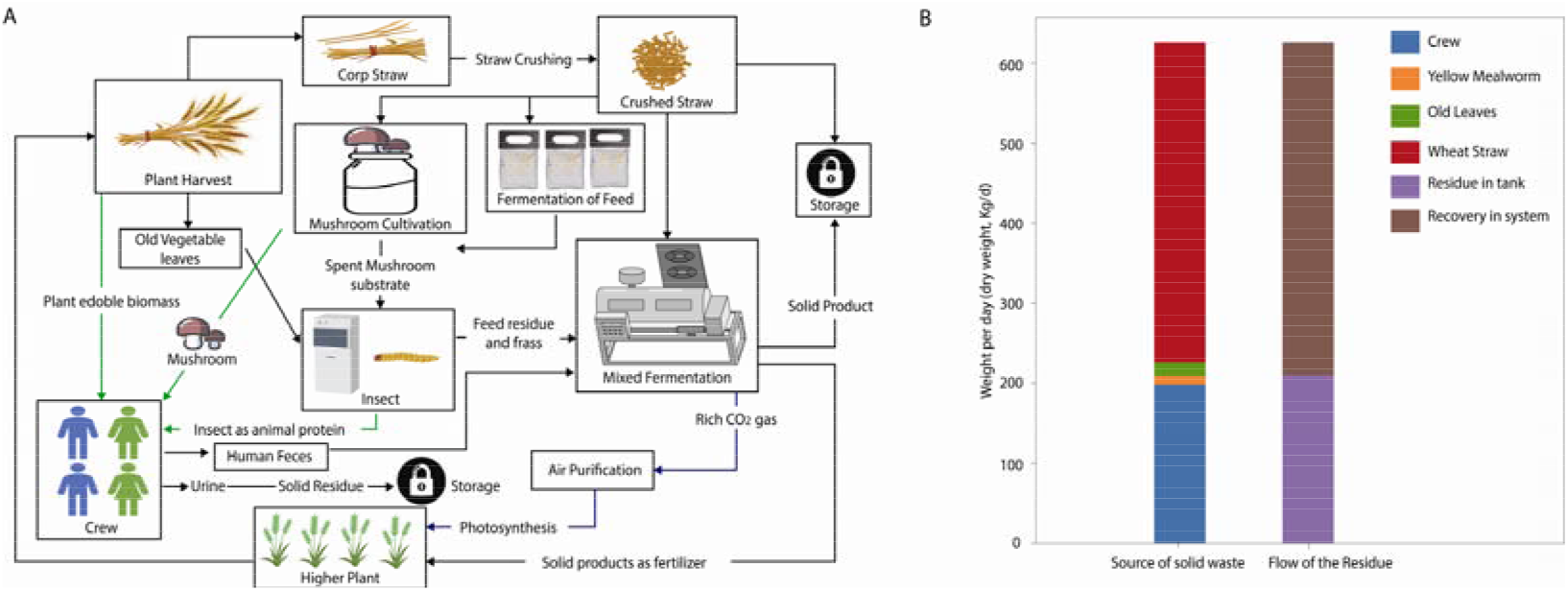
Solid waste processing. (A) Solid waste recycling processes in Lunar Palace 1. (B) The dry weight of solid waste in the “Lunar Palace 365” experiment.

### 2.3 Plant cultivation

The plant cabin was a critical component unit. Its internal cultivation device had a three-layer stereoscopic design, with a total planting area of 120 m^2^. Considering human energy and nutrition requirements^22^, nutrition compositions of plant-based food^23^ and the diversity and richness of food, 35 kinds of plants (including five kinds of food crops, 29 kinds of vegetables and one kind of fruit) were selected, and the corresponding planting zoning design was determined (Table S1).

Based on the light requirement of each plant^21^ and our previous studies^2^, the planting were assigned to the two plant cabins with different light parameters. For plant cabin I, the lighting was continuous (24/0 h light/dark), with the light intensity of 300-800 μmol m^−2^ s^−1^. For plant cabin II, the light cycle was 12/12 h light/dark with light from 08:00 to 20:00 everyday and a light intensity of 250-800 μmol m^−2^ s^−1^. Vermiculite or soil-like substrate converted from solid wastes was used for solid media culture in plant cabin I, and hydroponics were used for plant cabin II. Hoagland nutrient solution was used for irrigation in both plant cabins.

### 2.4 Experimental design and crew selection

We also investigated the impact of scenarios that might happen during actual lunar missions, including crew shift changes and simulated accidents, on the stability of the system. Further, effects of different lighting environments (i.e., whether the window was blocked or not) on the crew’s emotions and biological rhythms were explored. The timeline of “Lunar Palace 365” experiment is presented in Fig. S1. The experiment has three phases: 1) Group I was on duty at the initial phase of 60 days; 2) Group II then stayed instead in the system for a record-breaking 200 days for the second phase; 3) Group I re-entered the cabin to replace Group II for the 105-day third phase, with an additional “accidental” five-day extension for the psychological testing of the crew members. In the last month, three simulated power loss accidents for two, three and four hours were conducted. The purpose of the accident scenarios was to assess both the changes in CO_2_ concentration in the system when the power was accidentally down and the time for the system to recover to a stable state.

There were crew shift changes between each two adjacent phases of the experiment, which took place on the 60^th^ and the 260^th^ days. In order to prevent any gas exchange between the system and the outside during shift changes, the crew was required to enter the cabin through an airlock chamber (Fig. S2). For each shift change, the incoming crew would first enter the cabin through the airlock chamber at the time of 10:00 and conduct mission handover work in the cabin with the outgoing crew. After seven hours (at 17:00), the first set of volunteers would leave the cabin through the airlock chamber to complete the shift change. During the shift change process, there were a total of eight crew members in the cabin for a long duration of seven hours, serving as an overload disturbance to the system. Throughout the experiment, concentration changes of O_2_ and CO_2_ in the system were measured by sensors (FGD10B, Status Scientific Controls Ltd, Status Scientific Controls Ltd, UK) and logged by the control system.

Based on the experimental design, the eight volunteers chosen as crew members of “Lunar Palace 365” were divided into two groups (4 volunteers/each group, including two males and two females). All volunteers were aged 24 to 30 years old and tested as physically and psychologically healthy, without unhealthy habits or serious diseases. Basic characteristics of the volunteers are presented in Table S2. The respiration-CO_2_ production rate after moderate labor of crew group II was about 10% higher than that of crew group I (Fig. S3), suggesting a difference in the metabolic rate of the two crew groups. The labor, including plant cultivation unit maintenance, waste treatment unit maintenance, food harvest and food processing, health index detection etc., were clearly assigned to the four members in each group. Daily working and sleeping hours were both eight hours.

## 3 Results and discussion

### 3.1 Atmosphere control

#### 3.1.1 O_2_ and CO_2_ balance

Maintaining O_2_ and CO_2_ concentrations in an appropriate range is a basic indicator of BLSS stability. Concentrations of CO_2_ and O_2_ in the cabin throughout the experiment are shown in Fig. 4A. In the dark phases from 20:00 to 08:00 at the next day in plant cabin II, this cabin was not a producer of O_2_ but a consumer, and thus the CO_2_ and O_2_ concentrations showed daily fluctuations. Overall, the CO_2_ concentration was held between 246–4,131 ppm throughout the experiment, with an average CO_2_ concentration of 1241 ppm. Accordingly, O_2_ concentration in the system was maintained within 20.11%–21.52%, with an average of 20.71%. All these data are very close to the design standards of NASA’s latest advanced life support system for the atmospheric environment level of crew requirements (21.2% oxygen) and plant requirements (1300 ppm CO_2_)^24^. In general, for BLSSs, and similar compact closed spaces such as the international space station (ISS), it is essential to avoid possible accumulation of CO_2_ so as to avoid negative effects on humans and plants. Recent NASA studies have shown that excessive CO_2_ could lead to headaches and reduce cognitive function of the crew^25,26^. Furthermore, NASA reduced the maximum allowable CO_2_ concentration of ISS to 4 mmHg (5,332 ppm), with an evidence that CO_2_ concentration between 0.5 mmHg (666 ppm) and 2.0 mmHg (2,667 ppm) is optimal to maintain the health and performance of astronauts^26,27^. Our prior studies also demonstrated that, although an appropriate increase in atmospheric CO_2_ content increased plant yield, this beneficial effect disappeared when CO_2_ was over 2,000 ppm, while an inhibitory effect to the plant occurred when CO_2_ levels exceeded 5,000 ppm^28,29^. Compared with the BIOS-3 experiment^3^ of Russia and the 105-day Lunar Palace 1 experiment^2^, dynamic atmospheric data of our current system showed a higher stability in CO_2_ and O_2_ concentrations.

**Fig.4.**
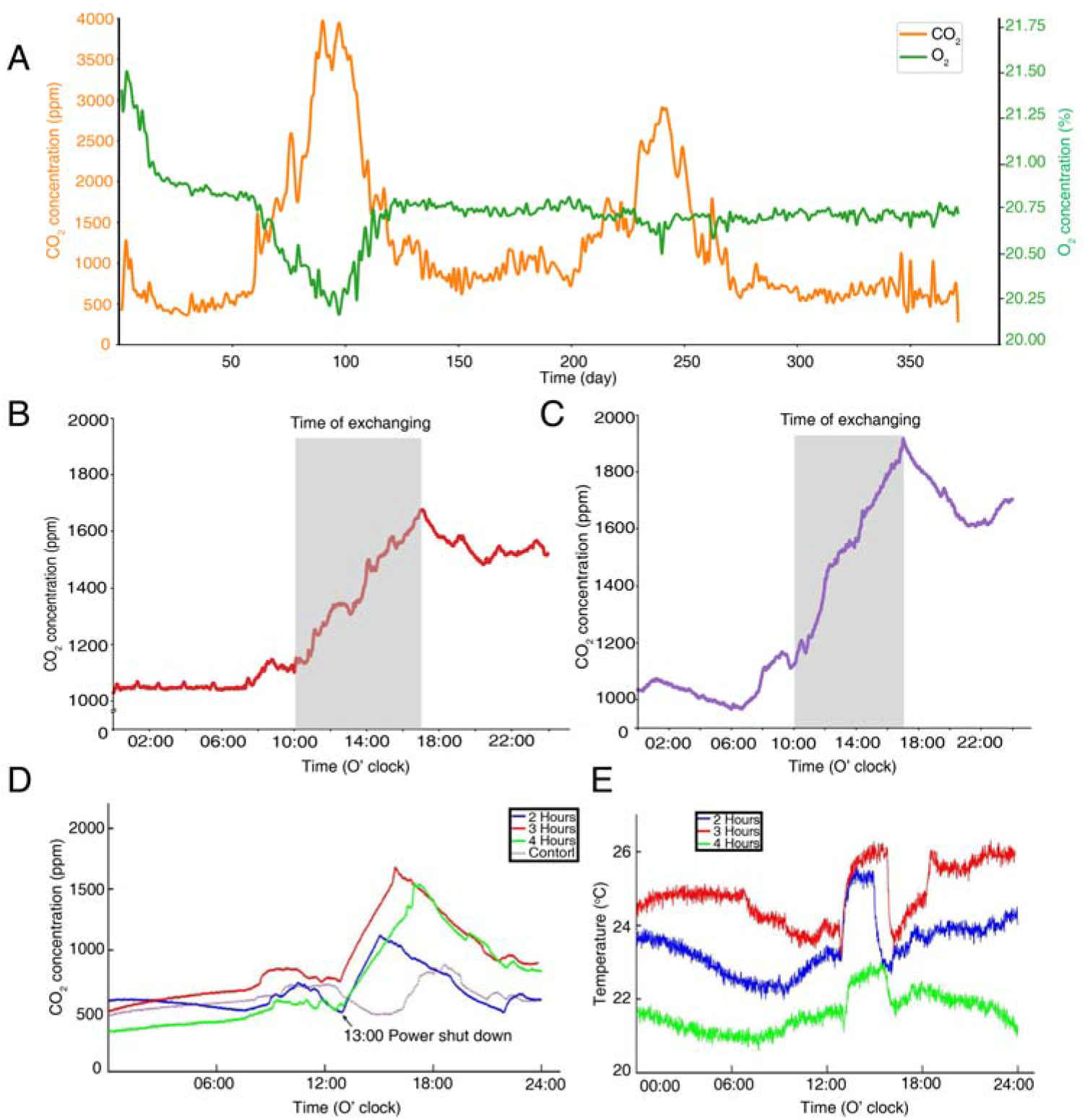
O_2_ and CO_2_ fluctuations. (A) The levels of CO_2_ and O_2_ over the course of the entire “Lunar Palace 365” experiment (370 days). (B) The change in carbon dioxide concentration around the first crew shift change (28h). (C) The change in carbon dioxide concentration around the second crew shift change (24h). (D) The changes in CO_2_ concentration in the simulated power loss experiments in the Lunar Palace 1. (E) The changes in temperature in the simulated power loss experiments in the Lunar Palace 1.

In addition to the short-term peak of CO_2_ gas resulting from the crew shift changes (the 60^th^ and 260^th^ days), there were two longer peaks of CO_2_ gas fluctuation (61–120 days and 200–260 days), both falling in the second phase of the experiment (group II) (Fig. 4A). Two reasons were attributed to the first long peak (after the 60^th^ day): 1) Unlike rigid physical-chemical systems, Lunar Palace 1 was a biological system with stronger adaptability to changes. Immediately after the crew shift change, the incoming group II had a higher metabolic rate that generated a higher-load impact on the system. Adaptation of the biological system was a slow process, leading to the accumulation of CO_2_ right after the shift change. 2) When the CO_2_ exceeded 3,000 ppm, it inhibited plant growth. This further aggravated the accumulation of CO_2_. The second long peak (after the 260^th^ day) could be attributed the artificial inhibition of wheat growth during the test of the resilience and recovery function of the biological system. Plant unit was the only unit in the system able to regenerate O_2_, in which wheat occupied the largest planting area in Lunar Palace 1. Therefore, inhibition of wheat growth could have a large impact on the gas stability of the system. The system atmosphere returned to a relatively stable state after two CO_2_ fluctuation peaks by adjusting photoperiods of plant units^2,18,30^, fermentation rates of waste units^31^ and other biological regulation factors. All these findings verify the elasticity of the biological system and the effectiveness of biological units in regulating atmospheric balance. In addition, the elevated CO_2_ and the corresponding O_2_ decreases also verify the airtightness of Lunar Palace 1 system, as shown in Fig. 4A. During shift changes, carbon dioxide rose from its initial stable state and then fell slightly before settling into a new stable state (Fig. 4B & Fig. 4C), indicating that the system was highly regulated.

When the BLSS power was “accidentally” lost, the plants were unable to photosynthesize due to the dark, while humans and other animals continued to produce CO_2_. Thus, the concentration of CO_2_ in the system increased (Fig. 4D). When the initial CO_2_ concentration in the system was lower than 600 ppm, the power loss resulted in an increase of CO_2_ concentration, which remained below 1,800 ppm at a rate of 251.0 - 312.3 ppm/h within four hours. When the power was restored, CO_2_ returned to a stable state within 12 - 24 hours (Fig. 4D), indicating that atmospheric fluctuations caused by power loss was small enough for the system to quickly adjust. It is possible that the increase in CO_2_ (to ~2,000 ppm) improved plant photosynthesis after the power was restored^28,29^, leading to a new CO_2_ concentration stable state in the system in a short time (Fig. ???). Moreover, increasing CO_2_ production rate was closely related to the temperature of the system (Fig. 4D & Fig. 4E), which is consistent with a previous study^32^ pointing out that plant respiration intensity positively correlates with system temperature within a certain temperature range. Importantly, a 100% recycling rate of O_2_ was achieved without any external gas input.

#### 3.1.2 Trace harmful gases

During the 370-day airtight experiment, 12 trace harmful gases, including hydrogen sulfide, were tested periodically. Concentrations of these gases were all controlled within a certain range (Fig. S4). We compared these data with the allowable concentration of air composition in a Chinese nuclear submarine cabin (GJB11A-98), the maximum allowable concentration of harmful gases in NASA manned spacecraft (JSC20584), and the maximum allowable concentration of harmful gas in Russian manned spacecraft (ГOCT Р50804-95) (Table S3). We found that the highest concentrations of harmful gases in our experiment were far below the maximum allowable concentrations of the Chinese standards, and only the concentrations of acrolein and ozone were slightly higher than those allowed by NASA and Russia^33–35^. We expect that acrolein was a trace gas readily produced by the decomposition of food and condiments in the cooking process, while the trace ozone may have been produced by the ultraviolet light source. Taken together, these results indicate that the BLSS can effectively control the concentrations of trace harmful gases by air purification of the plant unit, thus providing the crew with a healthy living and working environment. These results also support previous findings that there was no continuous accumulation of trace harmful gases in closed ecosystems due to the absorption and removal of biological units^3^.

### 3.2 Food production

Table S4 shows the production efficiency of crops during the “Lunar Palace 365” experiment. Crop yields were all greater compared with those in the “Lunar Palace 105” experiment^2^. However, compared with the BIOS-3 system in Russia^36^ and the NASA basic value of advanced life support system in the United States (ALS-BVAD)^21^, some crops in our experiment exhibited lower yields. The reason might be that there are differences in crop varieties, which means that more productive crop varieties that are suitable for growing in closed artificial ecosystem need to be developed. The food regeneration rate in the system, calculated in dry weight, was 73% throughout the experiment (Fig. S5). In fresh weight, the regeneration rate was 83% instead, with a 100% plant-based food regeneration rate. The food regeneration rate in dry weight (73%) in the current four-person 370-day experiment was higher than that in a previous three-person 105-day closed experiment (55%)^2^, a four-person 180-day closed experiment (55%)^19^, and the three-person 90-day BIOS-3 experiment in Russia (48%)^3^, but lower than that in the Japanese two-person 28-day CEEF system experiment (92% - 95%)^37^. It is important to note that the food regeneration capacity of a system not only depends on the crew size, but also relies on the planting area size of crops. Compared with the larger planting area in the other systems, such as the four-person, 180-day closed experiment system (260 m^2^) and the Japanese two-person 28-day CEEF system (150 m^2^), the planting area of our present experiment was only 120 m^2^. Therefore, our experiment exhibited a relatively lower resource consumption.

### 3.3 Water recovery

Fig. 2B shows the details of the water utilization and regeneration in the experimental system of “Lunar Palace 365”. Water consumption of each crew group was 74.37 kg/d, and daily water consumption of each crew member was 18.59 kg person^−1^ day^−1^. These values fall between the designed water demand of an early planet base and a mature planet base^21^. Total transpiration of the plant cabin was 724.26 L/d, and the average transpiration rate over the 120 m^2^ planting area was 6.13 L m^−2^ day^−1^, which is similar to that obtained in previous reports^5,38^. Condensed water generated by the system after purification and boiling was in line with the drinking water standards of China, the United States, and Russia (Table S5). Reclaimed water from wastewater purification conformed to the standards for hydroponics water for agricultural irrigation (Table S6) and was directly used as nutrient solution for plant irrigation. There was no external water input during the entire experiment, with a 100% recycling of water.

### 3.4 Solid waste bioconversion

The logistics of waste treatment and recycling in the “Lunar Palace 365” experiment is presented in Fig. 5. In this experiment, urine and fresh feces produced by each crew group per day were 7,130 g and 779 g (about 198 g of dry weight), respectively. Daily metabolic waste per person was 1,783 g of urine and 195 g of fresh feces. The mass of urine and feces varies with the quantity and composition of consumed food and drinking water^21^. The urine amount in the current experiment were within the range observed by NASA (1,390-2,107 g person^−1^ day^−1^) in previous studies^21^, but the feces were slightly more than those measured by NASA (20.5-132.0 g person^−1^ day^−1^). This may be because the diet of the crew in the current experiment had more abundant fiber and additional fiber in the diet was known to increase daily stool mass^39^.

**Fig.5.**
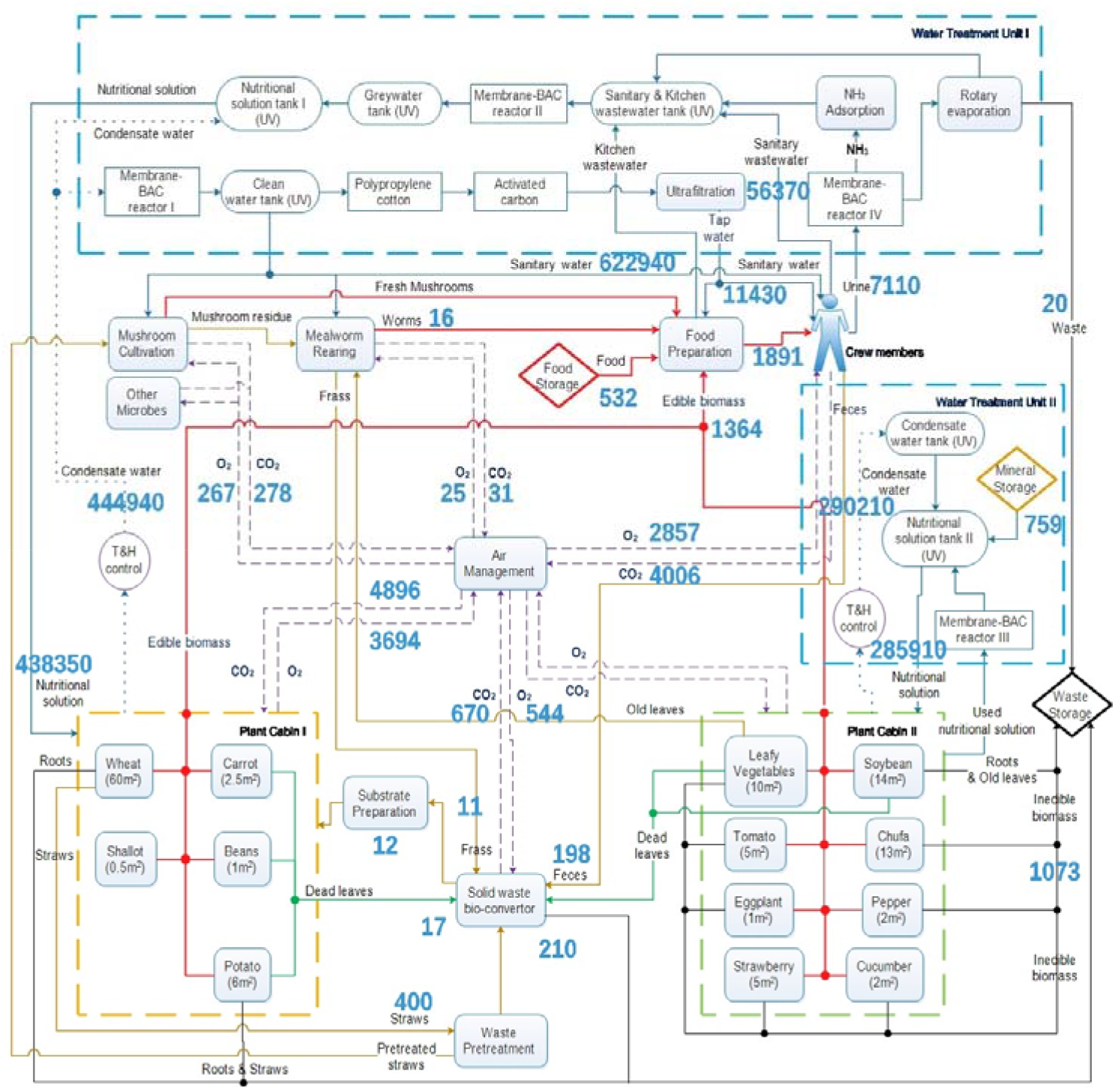
Mass flow relationships among system components in the “Lunar Palace 365” experiment (g/d). Arrows indicate the flow direction of mass. Object boxes represent each biological unit. The yellow, light green and blue dotted frames represent plant cabin I, plant cabin II and water treatment units. Storage waste matter, liquid, gas and food flows are marked by black, blue, purple and red arrows, respectively. The green and yellow arrows represent the cycle of solid matter. Data of solid matter was on a dry-weight basis.

The urine was distilled under low pressure after biological pretreatment, recovering 7,110 g water with a 99.7% recovery of nitrogen (Fig. 5). The residual urine slag was stored (Fig. 2A). The recovered water was further treated to prepare nutrient solutions for plant irrigation (Fig. 2A). Compared with the previous 105-day “Lunar Palace 105” experiment^2^, the ability of the current system to treat the urine increased by 1.22-fold (increased from 3,295 to 7,130 g/d). This ensured the requirement of irrigation water for plants and further ensured the amount of drinking water for crew.

Feces produced daily by the crew, together with non-edible biomass of plants and yellow mealworm feed residue (vegetable old leaf and insect dung) were fermented. The total, cumulative dry weights of straw, human feces, and yellow mealworm feed residue were 151,129 g (including the initial 3,000 g), 73,233 g, and 10,403 g, respectively, during the 370-day experiment. At the end of the experiment, the dry weight of remaining fermentation material in the solid waste fermentation device was 7,826 g. Therefore, the waste degradation rate of the solid waste fermentation unit was 67%, with the microbes degrading 400 g of dry solid waste on average per day. The key role of microbial degradation in the waste treatment was to make element carbon in the waste enter the atmosphere to rejoin the material cycle. Based on changes in carbon content before and after fermentation, the solid waste fermentation device contributed 670 g of CO_2_ to the atmosphere per day, being an essential part of the material cycle. The solid waste degradation rate of 67% in the current experiment was a significant improvement upon the rate of 41% in the “Lunar Palace 105” experiment also taken place in the facility of Lunar Palace 1^2^. Additionally, daily solid waste degraded by microorganisms was increased by 259 g compared to the “Lunar Palace 105” experiment. This demonstrates the performance of the solid waste treatment unit in the current Lunar Palace 1 system. It is worth noting that oxygen consumption of exogenous input must be considered in the calculation of solid waste treatment capacity of the system. To maintain the mass were balanced and stable in BLSS with exogenous input food and condiments for crew, and the equal amount of waste needed to be exported outside the system, which can prevent the accumulation of elements and substances that can cause the system to become unbalanced. Hence, the system optimization should not be based solely on improving the recycling rate of waste such as inedible plant parts and passengers’ excrement^2^. The design of each unit of BLSS and the setting of parameters must be considered from the perspective of the whole system, so that each biological unit in the system and the cabin environment can match each other, and the scientific rationality of the overall design cannot be ignored only in pursuit of individual technical indicators.

### 3.5 Energy expenditure and health status of the crew

Through accelerometers worn by crew members, daily energy consumption of each member was continuously monitored during the experiment (Fig. S6). Average energy consumption was 2,200 to 2,600 kcal per day for males, and 1,400 to 1,800 kcal for females. These data are similar to that obtained in the 105-day three-person “Lunar Palace 105” experiment^2^. However, the energy consumption of males was slightly higher than that in the CEEF experiment in Japan (1900-2300 kcal/d)^37^. Human basal energy consumption (BEE) had three components: resting metabolism (basal metabolic), physical activity, and the thermic effect of food^40^. In our study, the crew’s composition ratio of the three nutrient classes (protein: carbohydrate: fat = 15 : 6 : 25) was similar to Japanese CEEF, indicating no significant difference in the thermal effect of food between the two experiments. Therefore, the difference in the energy consumption between Lunar Palace 365 and CEEF were likely due to differences in crew’s resting metabolism and physical activity. Moreover, the median energy consumption per person in group I and group II were about 7500 and 8400 kcal, respectively. The total energy consumption of crew group II was about 112% of that of group I, which also confirmed the correctness of the difference in metabolic levels between the two groups designed in this experiment.

In order to ensure the energy supply to the crew, we provided food supplies of 7800 kcal and 8700 kcal per day to groups I and II, respectively, throughout the experiment. The crew’s mental and physical health indices did not fluctuate significantly. Also, according to the physical examination results before and after the experiment, all crew members maintained a healthy status^41^.

### 3.6 System material closure coefficient

The proportion of recycled substances in the total substances is presented in Fig. 6. Based on the formula of “closure”^2,3^, the “Lunar Palace 365” experiment achieved a 100% recycling regeneration of O_2_ and water for life support of the crew members, 99.7% recovery rate of urine, 67% recovery rate of solid waste, 73% of the dry weight regeneration of food, and 98.2% of the system closure degree. The closure of the system was dramatically improved in contrast to the 105-day experiment of Lunar Palace 1.

**Fig.6.**
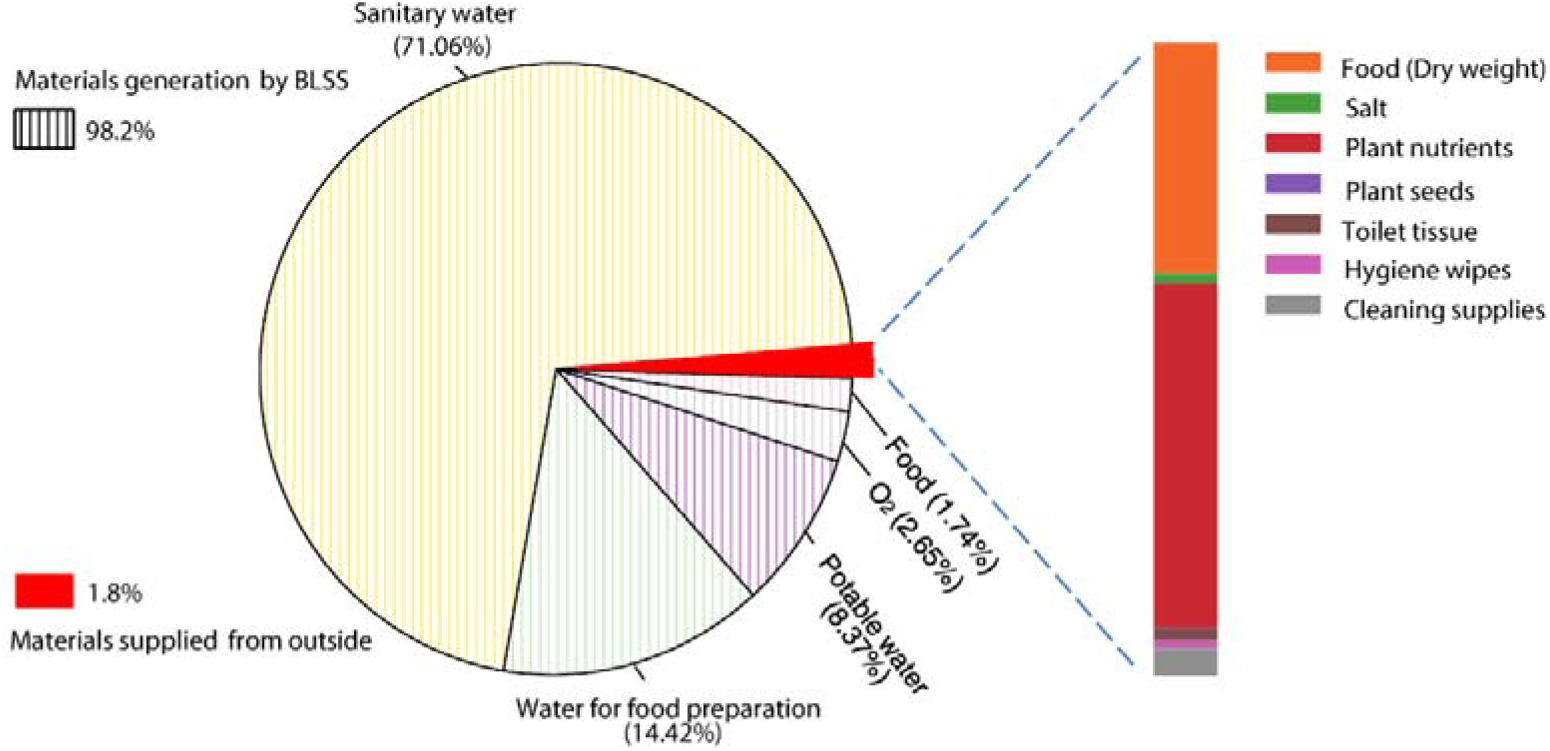
The proportion of recycled substances in the total substances.

## 4 Conclusion

The current study demonstrated that the atmosphere of the BLSS showed good stability under a multi-person load (4 persons) and long-term operation (370 days). The selected 35 plants grew well and exhibited high production efficiencies in the system. Water purification for personal hygiene and laundry reached the irrigation standard, and the purified potable water met the standards for condensate water. System closure degree was 98.2%. Those supported the feasibility and functional reliability of the Lunar Palace 1 system for long-term operation. Compared with our previous 105-day experiment, the current system demonstrated the ability to respond to increased CO_2_ caused by power loss and to quickly recover to a normal level. Moreover, disturbances in atmospheric CO_2_ concentration caused by shift changes was successfully eliminated by biological regulation, suggesting a good robustness of the system. The data also indicate that the advanced technology of a BLSS with an artificial, closed human-animal-plant-microorganism ecosystem had been achieved. For future work, quantitative research will be carried out in detail to better understand the impact of metabolic differences among crew members, shift change of members, and additional simulated accidents on the stability of the system. Computer simulated models should also be carried out using the data obtained from the current experiment to guide optimization of the BLSS system for practical application.

## Acknowledgments

This work was financially supported by the Pre-research Programme on Manned Spaceflight (PR China, No. 020302).

## Author contributions

Hong Liu and Yuming Fu designed the experiments. Zhihao Yi and Hui Liu performed the experiments. Yuming Fu, Zhihao Yi, Yao Du and Beizhen Xie analyzed the data. Yuming Fu, Zhihao Yi, Yao Du and Hong Liu wrote the manuscript. All authors reviewed the manuscript. All authors read and approved the final manuscript.

