## Supplementary materials for "Establishment of a closed artificial ecosystem to ensure human long-term survival on the moon"

***Corresponding author:**

Beihang University,

Phone numbers: +86 (010)82339837

Supplementary materials

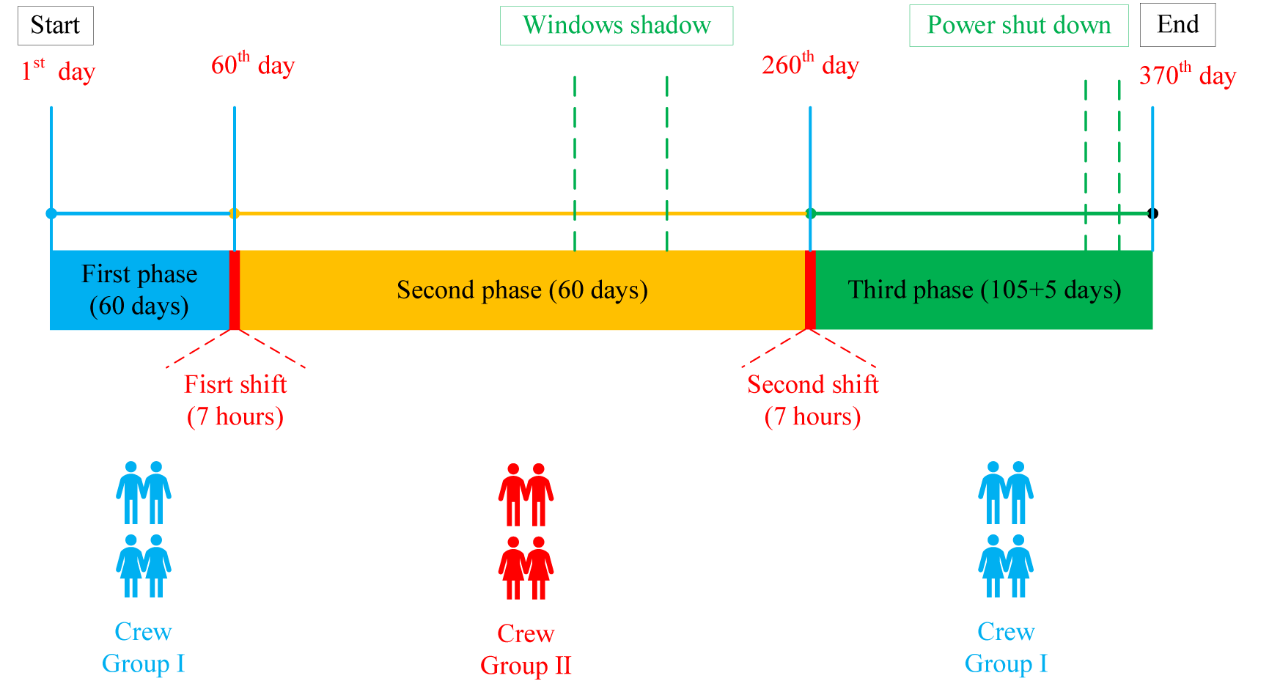

**Fig.S1 Timeline of the overall experimental design of the Lunar Palace 365 experiment.**

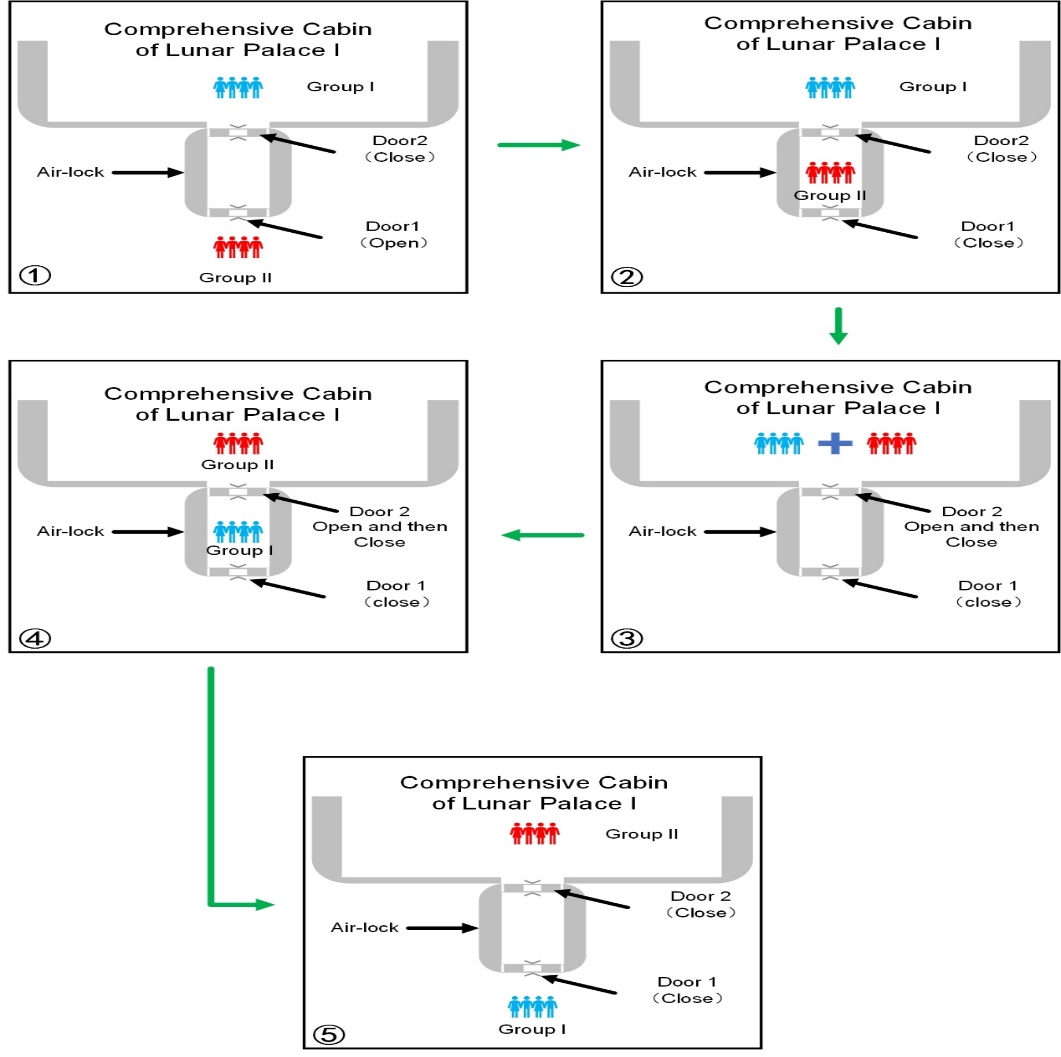

**Fig.S2 The change shifts of members with different metabolism states.**

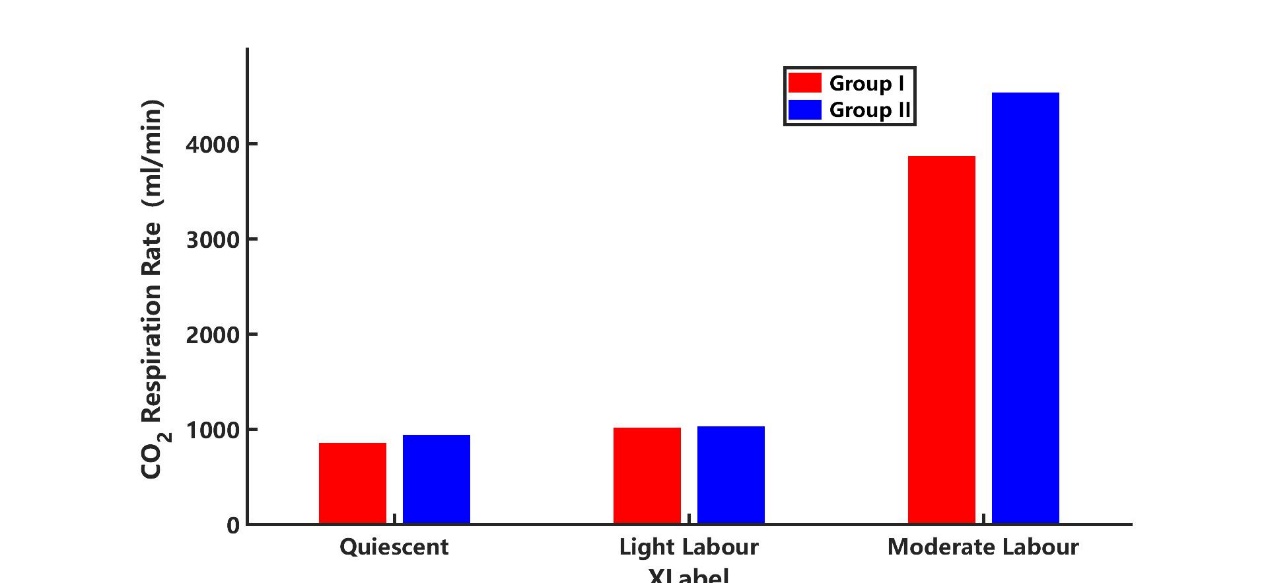

**Fig.S3 The rate of CO_2_ (produced by respiration) indicated the metabolic levels of the two crew groups in different metabolic states.** The metabolic levels of group II was about 10% higher than group I.

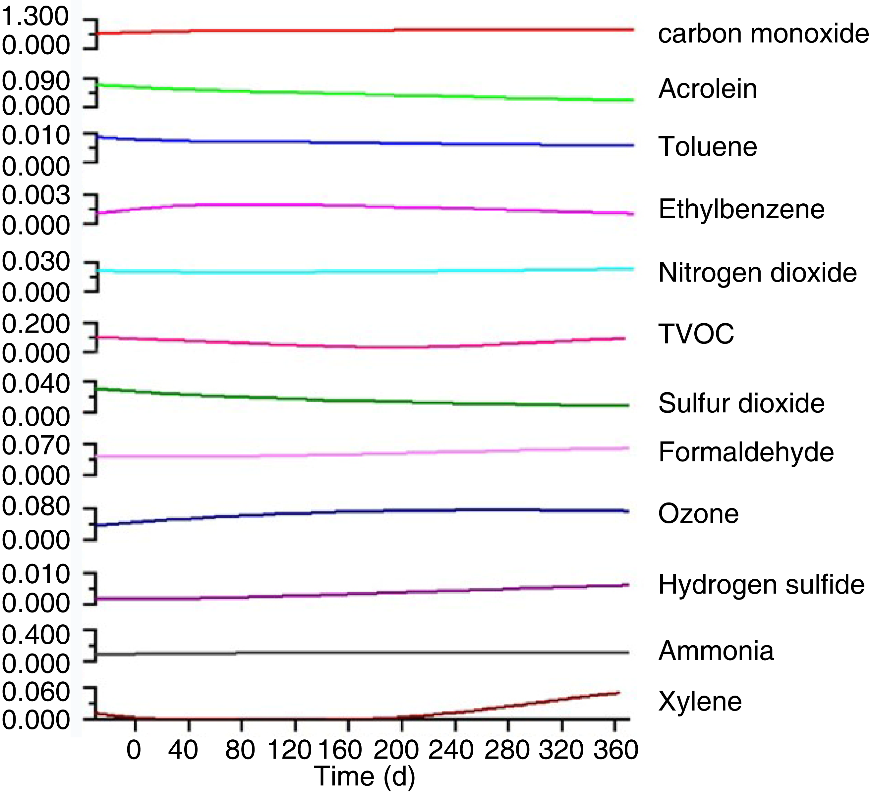

**Fig.S4 Contents of trace harmful gases (mg/m^3^) in the system during the Lunar Palace 365 experiment.**

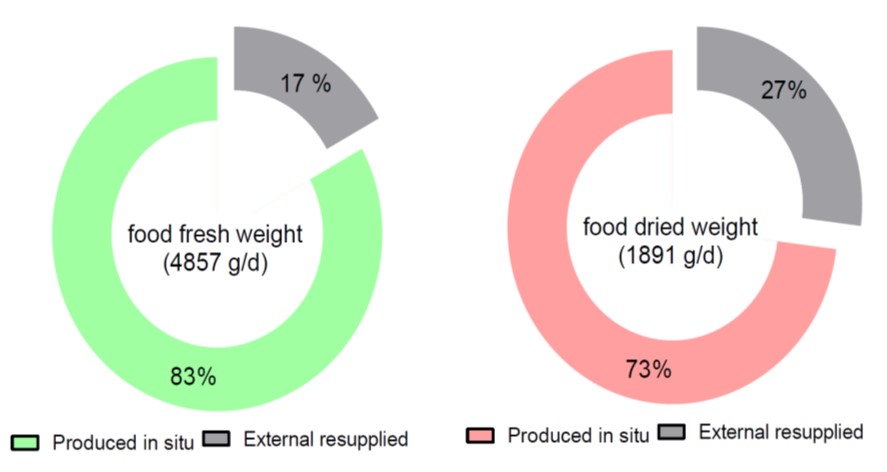

**Fig.S5 The plant food regeneration rate in the “Lunar Palace 365” experiment (left is fresh weight, right is dry weight).**

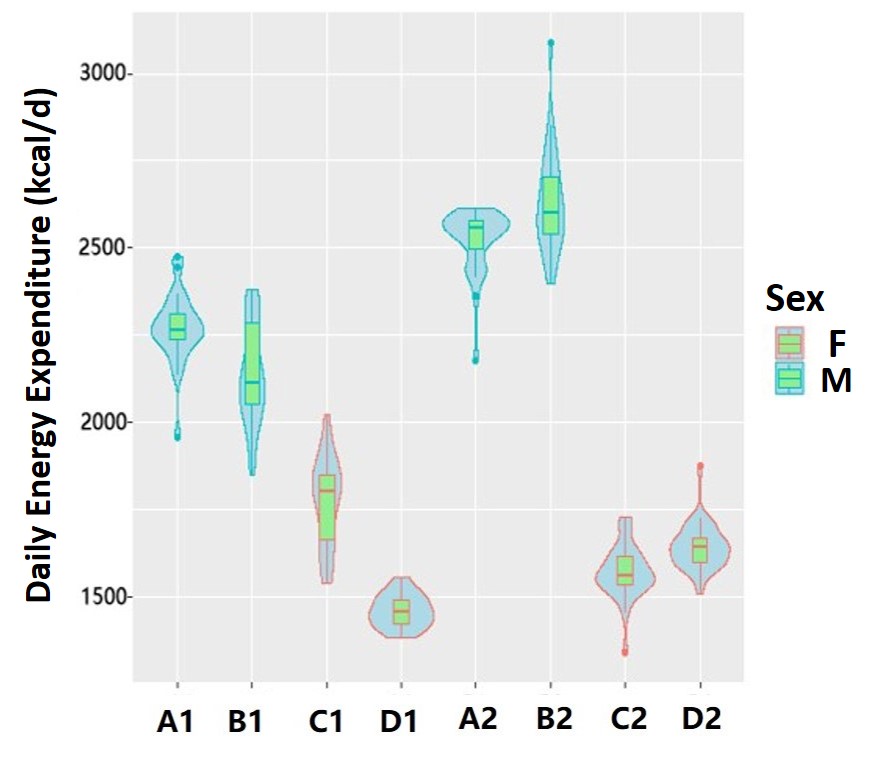

**Fig.S6 Daily energy expenditure of the crew volunteers (A1-D1, the first of group volunteers; A2-D2, the second group volunteers).**

**Table S1.** Design of plant species and acreage in the Lunar Palace 365 experiment

| Plant species | Life cycle（day） | Planting area （m^2^） | Time interval（day） | Location |
| --- | --- | --- | --- | --- |
| Wheat | 70 | 60-66 | 2-3 | Plant cabin I+ II |
| Carrot | 75 | 2.5 | 15 | Plant cabin I |
| Scallion | 42 | 0.5 | - | Plant cabin I |
| Potato | 120 | 6 | 10 | Plant cabin I |
| Kidney bean | 60-80 | 1.8 | 15-20 | Plant cabin I |
| Chufa | 133-144 | 6-13 | 10-24 | Plant cabin II |
| Soybean | 98 | 14 | 7 | Plant cabin II |
| Cucumber | 120 | 3 | 40 | Plant cabin II |
| Chili | 200 | 2 | 100 | Plant cabin II |
| Eggplant | 180 | 1-2 | 90 | Plant cabin II |
| Ornamental tomato | 180 | 3 | 60 | Plant cabin II |
| Infinite growth tomato | 240 | 1 | - | Plant cabin II |
| Leaf vegetables  (22 species) | 30-33 | 10-11 | 3 | Plant cabin II |
| Strawberry | 120 | 3 | 40 | Plant cabin II |

**Table. S2 Basic information on the Crew volunteers in the “Lunar Palace 365” ex**periment

| Group | Crew | Weight (kg) | Height (cm) | Age (year) | Gender |
| --- | --- | --- | --- | --- | --- |
| Group I | A1 | 64 | 180 | 24 | man |
|  | B1 | 63.5 | 168 | 30 | man |
|  | C1 | 56.5 | 165 | 29 | woman |
|  | D1 | 47 | 164 | 24 | woman |
| Group II | A2 | 66 | 174 | 27 | man |
|  | B2 | 72 | 185 | 26 | man |
|  | C2 | 52 | 157 | 25 | woman |
|  | D2 | 51 | 163 | 25 | woman |

Table. S3 Comparative analysis between maximum allowable concentrations in various countries and trace gas in the “Lunar Palace 365” experiment.

| Trace harmful gas | “Lunar Palace 365” experiment | Maximum allowable concentrations（mg/m^3^） | | |
| --- | --- | --- | --- | --- |
|  |  | China^[1]^  *(GJB11A-98)* | America^[2]^  *(JSC20584)* | Russia^[3]^  *ГОСТ Р 50804-95* |
| Carbon monoxide | 0.8~1.2 | 23 | 11 | 5 |
| Acrolein | 0.01~0.05 | 0.23 | 0.03 | - |
| Toluene | 0.01~0.51 | 40 | 60 | - |
| Ethylbenzene | <0.02 | 30 | 50 | - |
| Nitrogren dioxide | 0.019~0.027 | 0.8 | - | - |
| TVOC | 0.028~0.08 | 25 | - | - |
| Sulfur dioxide | 0.007~0.014 | 1 | - | - |
| Formaldehyde | 0.02~0.07 | 0.1 | 0.12 | 0.05 |
| Ozone | 0.06~0.076 | 0.08 | - | 0.03 |
| Hydrogen sulfide | 0.001~0.002 |  |  |  |
| Ammonia | 0.09~0.15 | 0.15 | - | 0.5 |
| Xylene | <0.06 | 10 | 6.5 | 5.0 |

[1]Fengtao G, Zhongquan L, Guogen C. Revision of Allowable Concentrations of Atmospheric Components in Nuclear Submarines (MAC GJB11-84). JOURNAL OF NAVY MEDICINE. 1999(1):7.

[2]James JT. Spacecraft maximum allowable concentrations for airborne contaminants. 2008.

[3]GOST R. 50804–95. Sreda obitaniya kosmonavta v pilotiruemom kosmicheskom apparate. Obshchie mediko tekhnicheskie trebovaniya [Cosmonauts habitable environments on board of manned spacecraft. General medicotechnical requirements]. Moskva. 1995. 71 p. Russ.

**Table. S4 Production efficiency of 13 selected crops (g/m^2^/d, dry weight) in exper**iments in different systems

| Crop | Lunar Palace 365 | Lunar Palace 105^[1]^ | BIOS-3^[2]^ | ALS-BVAD^[3]^ |
| --- | --- | --- | --- | --- |
| Wheat | 13.55±3.21 | 12.81±2.27 | 15.8 | 20 |
| Chufa | 10.78±4.96 | 6.87±3.12 | 26.04 |  |
| Soybean | 6.06±1.31 | 5.31±1.75 | - | 4.54 |
| Kidney bean | 10.18±1.36 | 10.34±1.09 | - | 11.88 |
| Leaf vegetables | 6.02±1.46 | 5.99±0.56 | 3.9 | 6.06-6.57 |
| Carrot | 17.90±6.36 | 3.58±0.21 | 20.4 | 8.98 |
| Cucumber | 6.97±1.64 | 10.03±1.46 | 15.5 | - |
| Scallion | 49.18 | 33.88 | 27.4 | 9 |
| Strawberry | 1.41 | 1.19 | - | 7.79 |
| Tomato | 18.86 | - | - | 23.17 |
| Potato | 6.05±2.46 | - | 4.8 | 30.08 |
| Chili | 17.15±2.97 | - | - | 23.17 |
| Eggplant | 8.76±2.33 | - | - | - |

[1]Fu, Y. et al. How to establish a Bioregenerative Life Support System for long-term crewed missions to the Moon or Mars. Astrobiology 16, 925-936 (2016).

[2]Gitelson, J. I. & Lisovsky, G. M. Man-made closed ecological systems. Vol. 9 (Crc Press, 2002).

[3]Hanford, A. J. Advanced life support baseline values and assumptions document. Technical Reports, 3 (2006).

Table. S5 Analysis of results of purification of condensate water (potable water and domestic water).

| Project | Unit | Condensate water | Maximum-contamination level in drinking-water | | | |
| --- | --- | --- | --- | --- | --- | --- |
|  |  |  | China^[1]^ | Russia^[2]^ | America^[2]^ | |
| pH | - | 7.34±0.43 | 6.5-8.5 | 5.5-9.0 | | 4.5-8.5 |
| Turbidity | NTU | <0.5 | 1 | 1.5 | | 1 |
| Chroma | Level | <5 | 15 | 20 | | n/a |
| COD_Mn_ | mg/L | 0.28 | 3 | - | | - |
| BOD_5_ | mg/L | <0.5 | - | - | | - |
| Nitrate-nitrogen (NO_3_-N) | mg/L | <3.25 | 10 | 10 | | 10 |
| Nitrite-nitrogen (NO_2_-N) | mg/L | <0.040 | 1 | - | | - |
| Ammonia- nitrogen  (NH_4_-N) | mg/L | 0.035 | 0.5 | 2 | | 0.5 |
| Arsenic | mg/L | <0.001 | 0.01 | 0.01 | | 0.01 |
| Chromium | mg/L | <0.004 | 0.05 | 0.1 | | 0.05 |
| Cadmium | mg/L | <0.0005 | 0.005 | 0.005 | | 0.005 |
| Lead | mg/L | <0.0025 | 0.01 | 0.05 | | 0.05 |
| Iron | mg/L | <0.0045 | 0.03 | 0.03 | | 0.03 |
| Aluminum | mg/L | <0.040 | 0.2 | - | | - |
| Mercury | mg/L | <0.0001 | 0.001 | 0.002 | | 0.002 |
| Cuprum | mg/L | <0.009 | 1 | 1 | | 1 |
| Cyanide | mg/L | <0.002 | 0.05 | 0.2 | | 0.2 |
| Phenol | mg/L | <0.002 | 0.002 | 1 | | - |
| Sulfide | mg/L | <0.02 | 0.02 | - | | 0.05 |
| Fluoride | mg/L | <0.01 | 1.0 | 1.5 | | - |
| Chloroform | mg/L | <0.00003 | 0.06 | - | | - |
| Carbon tetrachloride | mg/L | <0.00021 | 0.002 | - | | - |
| Total number of bacteria | CFU/mL | 91±100(0^*^) | 100 | 100 | | 100 |
| E. coli | CFU/100mL | 0 | 0 | <1 | | 0 |

[1] Hygienic standard for drinking water in chinese, GB5749-2006

[2] Advanced Life Support Requirements Document, JSC-38571C

Note：asterisk (*) represents the total number of bacteria in drinking water after heating

Table. S6 The quality of recycled water purified by biosorption technique in the “Lunar Palace 365” experiment.

| Project | Unit | Condensate water | Water Quality Standards of plant irrigation water in chinese^[1]^ |
| --- | --- | --- | --- |
| pH | pH units | 6.67±1.87 | 5.5-8.5 |
| COD | mg/L | 29.56±13.13 | 150 |
| TP | mg/L | 3.93±3.02 | - |
| Chromium | mg/L | <0.03 | 0.1 |
| Lead | mg/L | <0.001 | 0.2 |
| Zinc | mg/L | 0.42±0.17 | 2 |
| Copper | mg/L | 0.69±0.45 | 1 |
| Chlorine | mg/L | 6.47±2.86 | 350 |
| E. coli | CFU/100mL | 0 | 4000 |

[1] Water quality for agriculture in chinese, GB5084-2005
